# The 20-hydroxyecdysone agonist, halofenozide, promotes anti-*Plasmodium* immunity in *Anopheles gambiae* via the ecdysone receptor

**DOI:** 10.1101/2020.06.19.162081

**Authors:** Rebekah A. Reynolds, Hyeogsun Kwon, Thiago Luiz Alves e Silva, Janet Olivas, Joel Vega-Rodriguez, Ryan C. Smith

## Abstract

Mosquito physiology and immunity are integral determinants of malaria vector competence. This includes the principal role of hormonal signaling in *Anopheles gambiae* initiated shortly after blood-feeding, which stimulates immune induction and promotes vitellogenesis through the function of 20-hydroxyecdysone (20E). Previous studies demonstrated that manipulating 20E signaling through the direct injection of 20E or the application of a 20E agonist can significantly impact *Plasmodium* infection outcomes, reducing oocyst numbers and the potential for malaria transmission. In support of these findings, we demonstrate that a 20E agonist, halofenozide, is able to induce anti-*Plasmodium* immune responses that limit *Plasmodium* ookinetes. We demonstrate that halofenozide requires the function of ultraspiracle (USP), a component of the canonical heterodimeric ecdysone receptor, to induce malaria parasite killing responses. Additional experiments suggest that the effects of halofenozide treatment are temporal, such that its application only limits malaria parasites when applied prior to infection. Unlike 20E, halofenozide does not influence cellular immune function or AMP production. Together, our results further demonstrate the potential of targeting 20E signaling pathways to reduce malaria parasite infection in the mosquito vector and provide new insight into the mechanisms of halofenozide-mediated immune activation that differ from 20E.

## Introduction

Malaria kills over 400,000 people each year, with the majority of deaths in children under the age of five ^1^. Transmission requires an *Anopheles* mosquito to acquire and transmit *Plasmodium* parasites through the act of blood-feeding, a behavior that evolved in mosquitoes to acquire nutrient-rich blood for egg development ^2^. Strategies to interrupt mosquito blood-feeding, and subsequent parasite transmission, are an essential step in reducing disease transmission. The use of long-lasting insecticide treated bed nets (LLITNs) and indoor residual spraying (IRS) were integral in reducing malaria deaths by approximately 50 percent over the last 20 years ^1,3,4^. These preventative measures have proven effective in Africa, where over 90 percent of the malaria cases occur globally, yet progress in the remaining regions of the world have plateaued in recent years ^1^. Moreover, the increasing prevalence of insecticide and drug resistance now threaten a global resurgence in malaria cases ^1^, therefore requiring new approaches to combat malaria transmission.

In order to be transmitted, malaria parasites must survive several bottlenecks in their mosquito host, where parasite numbers at the oocyst stage are arguably the lowest in the malaria life cycle ^5^. These bottlenecks are mediated in part by blood-meal derived factors, the mosquito microbiota, and components of the innate immune system that ultimately determine vector competence and vectorial capacity ^5^. Therefore, targeting components of mosquito physiology and/or immunity can serve as excellent targets for malaria control strategies to interrupt *Plasmodium* development in the mosquito host.

Previous studies have demonstrated that the steroid hormone 20-hydroxyecdysone (20E) is essential for multiple aspects of mosquito physiology; such as development, mating, reproduction, and immunity ^6–9^. This includes the demonstration that 20E injection promotes *An. gambiae* cellular immunity and reduces both *E. coli* and *P. berghei* survival in the mosquito host ^8^. 20E agonists, which were initially developed as insecticides to reduce larval populations, have shown promise for use on adult *An. gambiae* mosquitoes ^10^. Methoxyfenozide (DBH) reduced mosquito lifespan, decreased fecundity, and significantly lowered the prevalence of *P. falciparum* parasite infection in the mosquito host ^10^. However, the mechanisms by which 20E agonists prime anti-*Plasmodium* immunity remain unexplored.

Here, we examine the 20E agonist, halofenozide, to better understand the mechanisms of anti-*Plasmodium* immunity stimulated by the direct application to the mosquito cuticle. In agreement with previous work on another 20E agonist ^10^, we demonstrate that halofenozide significantly reduces *P. berghei* parasite numbers when applied to mosquitoes prior to infection. Furthermore, we demonstrate that halofenozide treatment influences the success of *Plasmodium* ookinete invasion and requires the function of *ultraspiracle* (USP) as part of the heterodimeric ecdysone receptor (USP/EcR). When compared to earlier studies of 20E immune induction ^8^, we find that halofenozide and 20E differentially influence mosquito cellular immunity. These findings are supported by a recent study in *Drosophila*, which demonstrate differences in the physiological effects between 20E, a steroid hormone, and chromafenozide, a non-steroidal agonist ^12^. In summary, our study provides new detail into the mechanisms by which 20E agonists promote mosquito immunity and limit *Plasmodium* parasite survival.

## Results

### Halofenozide application reduces *P. berghei* survival and infection prevalence

Previous studies have demonstrated the ability of 20E and the 20E agonist, DBH, to induce mosquito responses that limit malaria parasite infection in *An. gambiae* ^8,10^. While recent evidence provides insight into the effects of 20E immune induction ^8^, the manner in which 20E agonists influence mosquito immunity have not been previously examined. To explore this question, we performed experiments with halofenozide, a similar 20E agonist, and examined its influence on malaria parasite infection. Halofenozide was topically applied to adult female mosquitoes without significant effect on mosquito survival (Figure S1) and challenged with *P. berghei* ~24hrs post-application. Compared to control (acetone-treated) mosquitoes, halofenozide treatment with either 0.25 μg/μL or 0.5 μg/μL significantly reduced the intensity of *P. berghei* oocysts eight days post infection (Figure 1A). Moreover, halofenozide reduced the prevalence of infection from 85% in control mosquitoes to <50% in halofenozide-treated mosquitoes (Figure 1B). Given the low mortality rate and the strong anti-*Plasmodium* effect, the remainder of the halofenozide experiments were conducted using the 0.5 μg/μL concentration.

**Figure 1.**
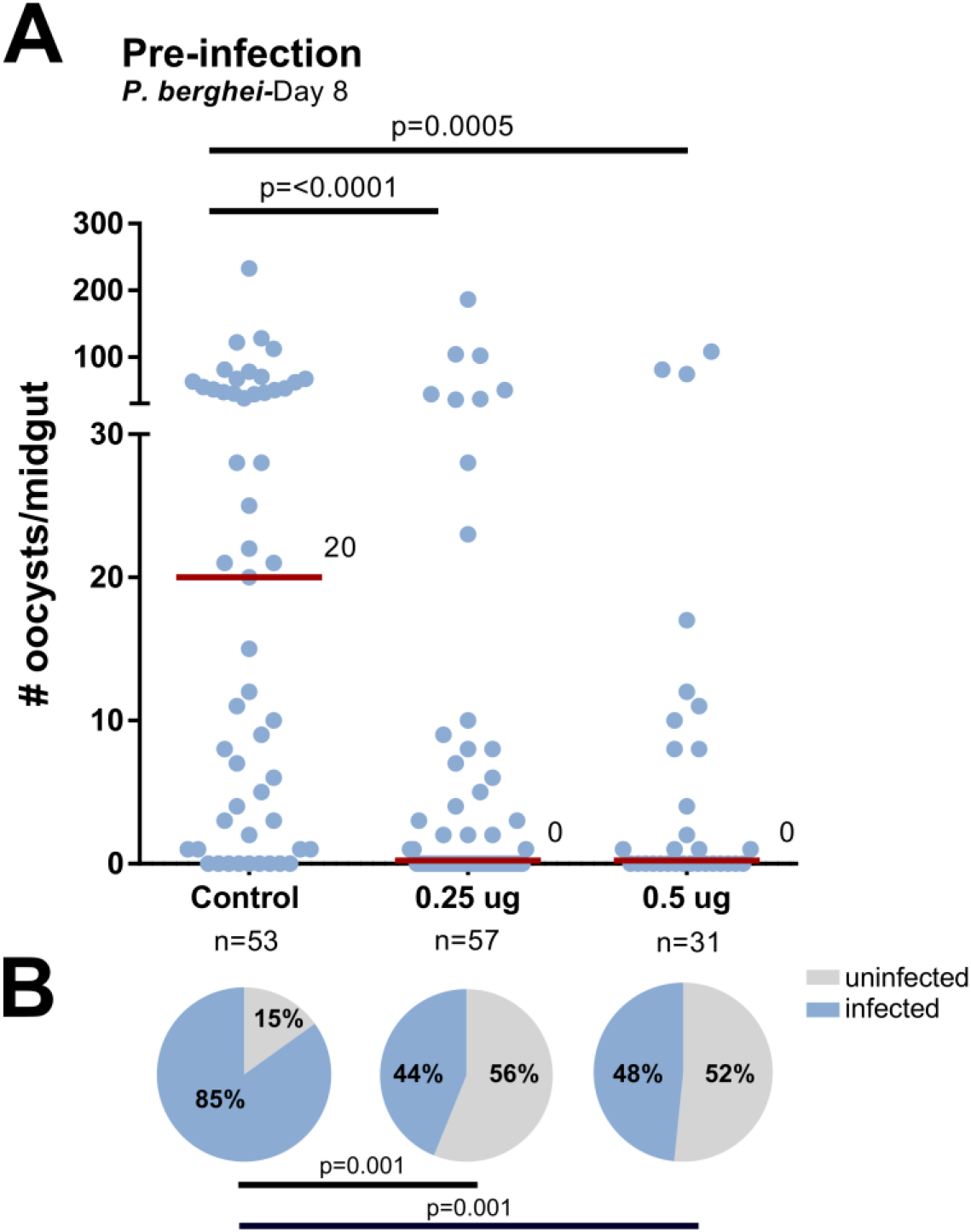
Halofenozide priming reduces *P. berghei* survival and infection prevalence. **(A)** 100% acetone (control) or halofenozide dissolved in 100% acetone (0.25 μg/μL and 0.5 μg/μL) were applied to *Anopheles gambiae* adult females 24 hours prior to challenge with a *P. berghei*-infected blood meal. Eight days post-infection, oocyst numbers were examined from dissected midguts. The red bar delineates the median number of oocysts from pooled data from three independent experiments. Data were analyzed by Kruskal-Wallis with a Dunn’s post-test using GraphPad Prism 6.0. n= the number of mosquitoes examined for each condition. **(B)** The prevalence of infection (% infected/total) is depicted for mosquitoes under each treatment and examined by Χ^2^ analysis to determine significance.

### Halofenozide promotes the killing of *P. berghei* ookinetes

To better determine how and when halofenozide limits parasite numbers, we examined the effects of halofenozide application on distinct stages of malaria parasite infection. When oocyst numbers were examined two days post-*P. berghei* infection, halofenozide-treated mosquitoes displayed a significant reduction in early oocyst survival (Figure 2A), suggesting that halofenozide application may promote ookinete killing. To further validate this point and determine if halofenozide application also influenced other stages of parasite development, halofenozide applications were performed approximately 24 hours post-*P. berghei* infection. Halofenozide treatment did not influence parasite survival when applied after an established *P. berghei* infection (Figure 2B), suggesting that the effects of halofenozide treatment are only effective against *Plasmodium* ookinetes when applied prior to infection.

**Figure 2.**
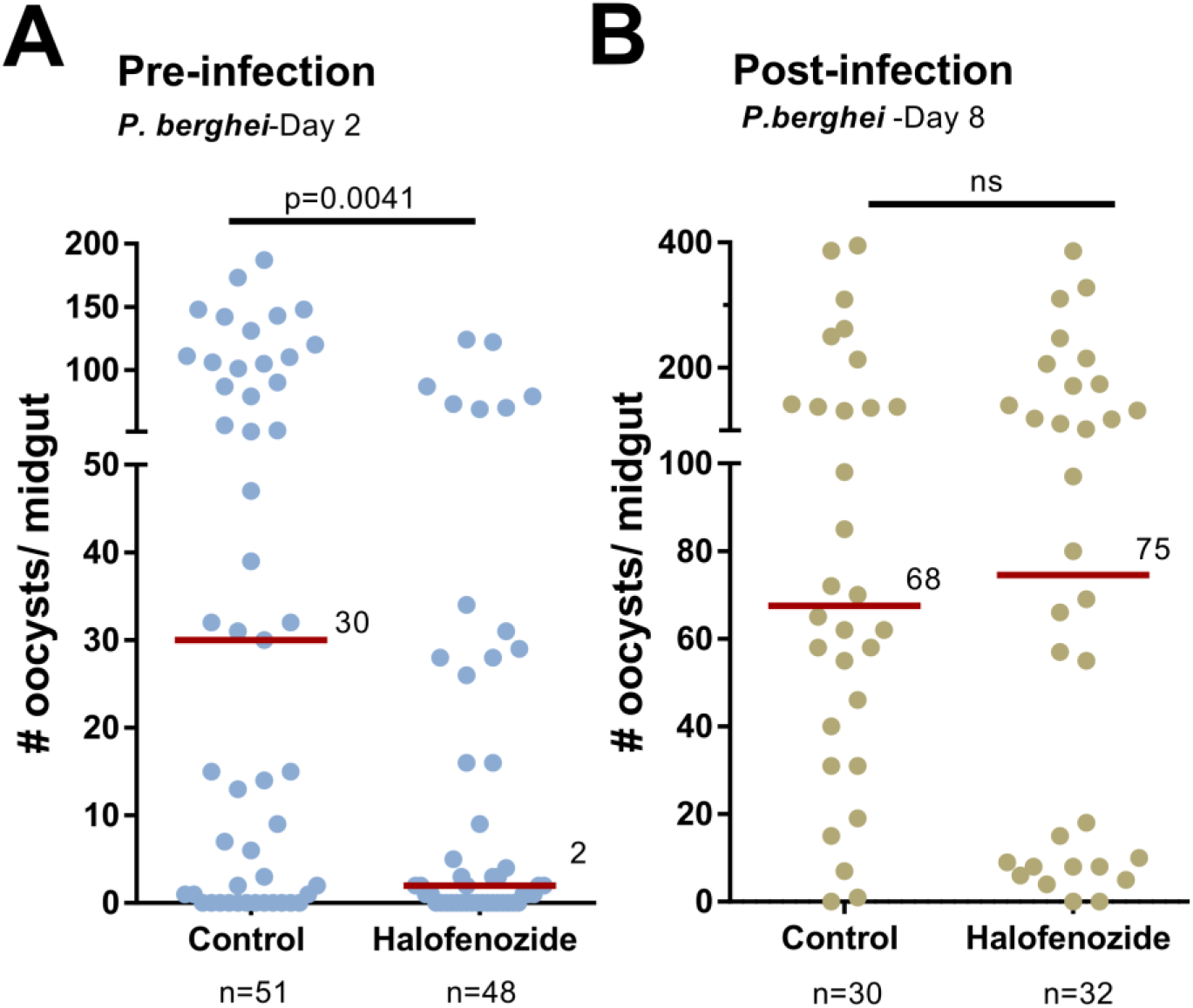
Temporal effects of halofenozide application on *Plasmodium* infection. The effects of halofenozide application were more closely examined to determine the influence of pre- **(A)** and post-infection **(B)** application on malaria parasite infection. **(A)** After priming with halofenozide (0.5 μg/μL), *P. berghei* oocyst numbers were examined two days post-infection. The significant influence of halofenozide on early oocyst numbers suggests that priming limits the success of ookinete invasion. This is supported by experiments where halofenozide (0.5 μg/μL) was applied ~24 hours post-*P. berghei* infection **(B)** to assess whether halofenozide impacts parasite survival if applied at a later time point. No differences in oocyst numbers were detected when oocyst number were evaluated eight days post-infection. The red bar delineates the median number of oocysts from pooled data from three independent experiments. Data were analyzed by Mann-Whitney analysis using GraphPad Prism 6.0. n= the number of mosquitoes examined for each condition.

To determine if halofenozide had an inhibitory effect on ookinete development or viability, we performed *in vitro P. berghei* ookinete cultures in the presence of 0.5 μg/μl of halofenozide. Treatment with halofenozide did not change the morphology or the number of ookinetes when compared to acetone controls (Figure 3A), supporting that halofenozide does not influence *Plasmodium* sexual stage development. Moreover, both control- and halofenozide-treated ookinetes were able to glide normally in Matrigel motility assays (Movie S1 and S2), suggesting that halofenozide treatment did not influence ookinete viability *in vitro*. Similar experiments were also performed *in vivo*, where *Plasmodium* sexual development was evaluated in the mosquito blood bolus 18 h after *P. berghei* infection. Fluorescent mCherry parasites were evaluated by morphology in control- and halofenozide-treated mosquitoes, displaying no differences in sexual stage development between treatments (Figure 3B). These data provide strong support that the reduced parasite numbers associated with halofenozide treatment are not the result of direct interactions on *Plasmodium* parasites.

**Figure 3.**
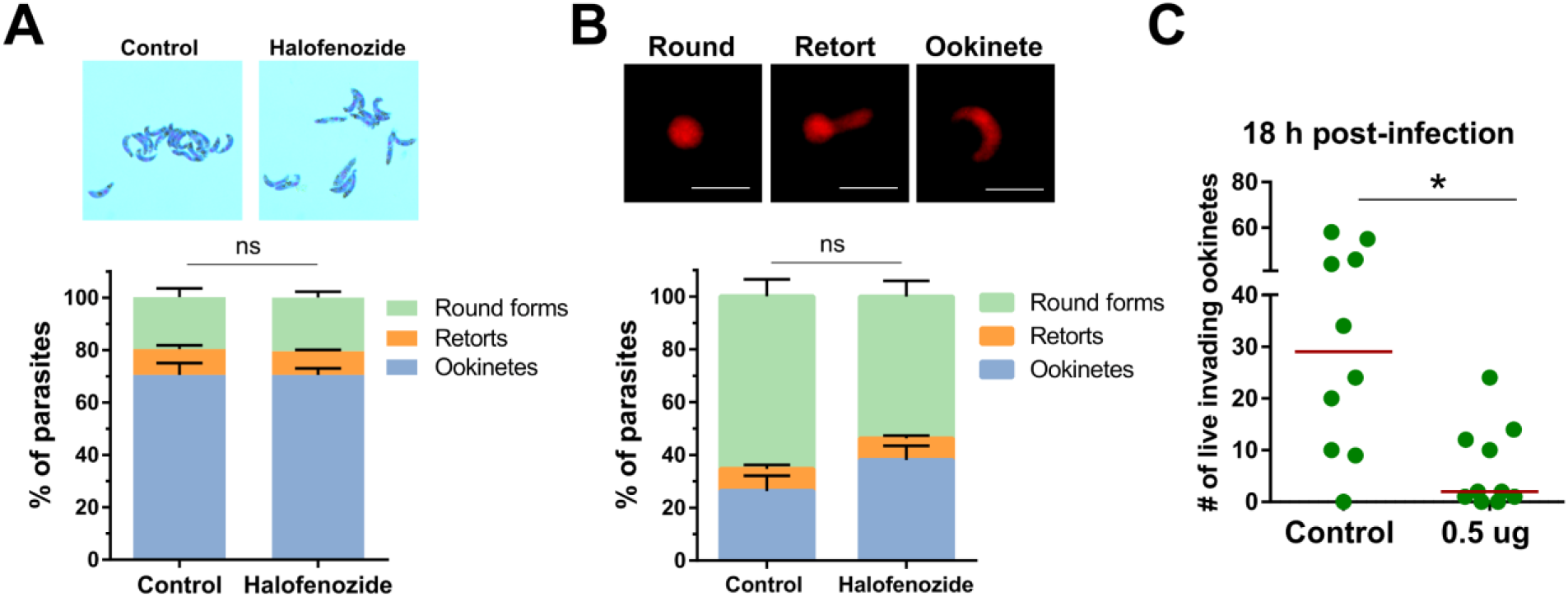
Halofenozide does not influence *Plasmodium* sexual development, but limits the success of ookinete invasion. The potential effects of halofenozide treatment on *Plasmodium* sexual development were examined through *in vitro* (**A**) and *in vivo* (**B**) experiments. When halofenozide was added to *P. berghei* ookinete cultures, ookinetes developed normally when examined by morphology (giemsa-stained images) with no differences in round forms, retorts, or ookinete numbers (**A**). Similar experiments were performed *in vivo*, in which the percentage of round, retort, and ookinetes were identified by morphology in mCherry parasites using fluorescence (**B**). For both **A** and **B**, data were collected from two independent experiments and were analyzed using a two-way ANOVA with a Sidak’s multiple comparison test. No significant differences were identified between treatments. ns, non-significant. (**C**) The number of live ookinetes (based on mCherry fluorescence) were examined in mosquito midguts samples 18h post-infection. Statistical significance was determined using Mann-Whitney analysis.

Since the mosquito gut microbiota are also major determinants of mosquito vector competence ^13,14^, we examined the potential that halofenozide could impact the microbiota. Levels of 16s rRNA expression, which serves as a proxy to assess bacteria numbers ^15,16^, were not significantly altered following halofenozide application (Figure S2A). Moreover, halofenozide had no impact on bacterial growth *in vitro* (Figure S2B), arguing that the anti-*Plasmodium* effects of halofenozide application do not involve alterations to the mosquito microbiota.

Additional experiments ookinete midgut invasion demonstrate that the number of live ookinetes (based on mCherry fluorescence) is significantly reduced following halofenozide treatment (Figure 3C). This suggests that the reduced numbers of early oocysts following halofenozide treatment (Figure 2A) are mediated by a yet undescribed mechanism of mosquito anti-*Plasmodium* immunity that limits the success of ookinete invasion. This also explains why halofenozide treatment after an established infection (Figure 2B; post-ookinete invasion) had no effect on parasite survival.

### Halofenozide requires USP to stimulate immune priming

To confirm the known function of halofenozide acting through the canonical 20E receptor ^17^, we examined the expression of *vitellogenin* and *cathepsin B,* genes responsive to 20E signaling ^18,19^. Halofenozide application significantly increased the expression of *vitellogenin* (Figure 4A; Figure S3) and *cathepsin B* (Figure 4B; Figure S3) to comparable levels as the injection of 20E (Figures 4A and 4B), supporting that halofenozide activates canonical 20E signaling. To determine if the heterodimeric ecdysone receptor comprised of the ecdysone receptor (EcR) and ultraspiracle (USP) is critical for the induction of anti-*Plasmodium* immunity following halofenozide treatment, we used RNAi to examine the role of the respective components in halofenozide immune activation (Figure S4). In RNAi experiments, the injection of dsRNA targeting the heterodimeric receptor had no effect on *EcR*, yet significantly depleted levels of *USP* (Figure S4). As a result, efforts to ascertain the function of the ecdysone receptor were evaluated with *USP*-silencing. In control, *dsGFP*-silenced mosquitoes, halofenozide application prior to *P. berghei* challenge resulted in a significant reduction in parasite numbers (Figure 4C), similar to previous results (Figures 1 and 2). However, the topical application of halofenozide in the *dsUSP-* silenced background (Figure S4) did not influence *P. berghei* survival (Figure 4C). Together, this suggests that halofenozide activation of anti-*Plasmodium* immunity requires the function of canonical 20E signaling through the heterodimeric ecdysone receptor (EcR/USP).

**Figure 4.**
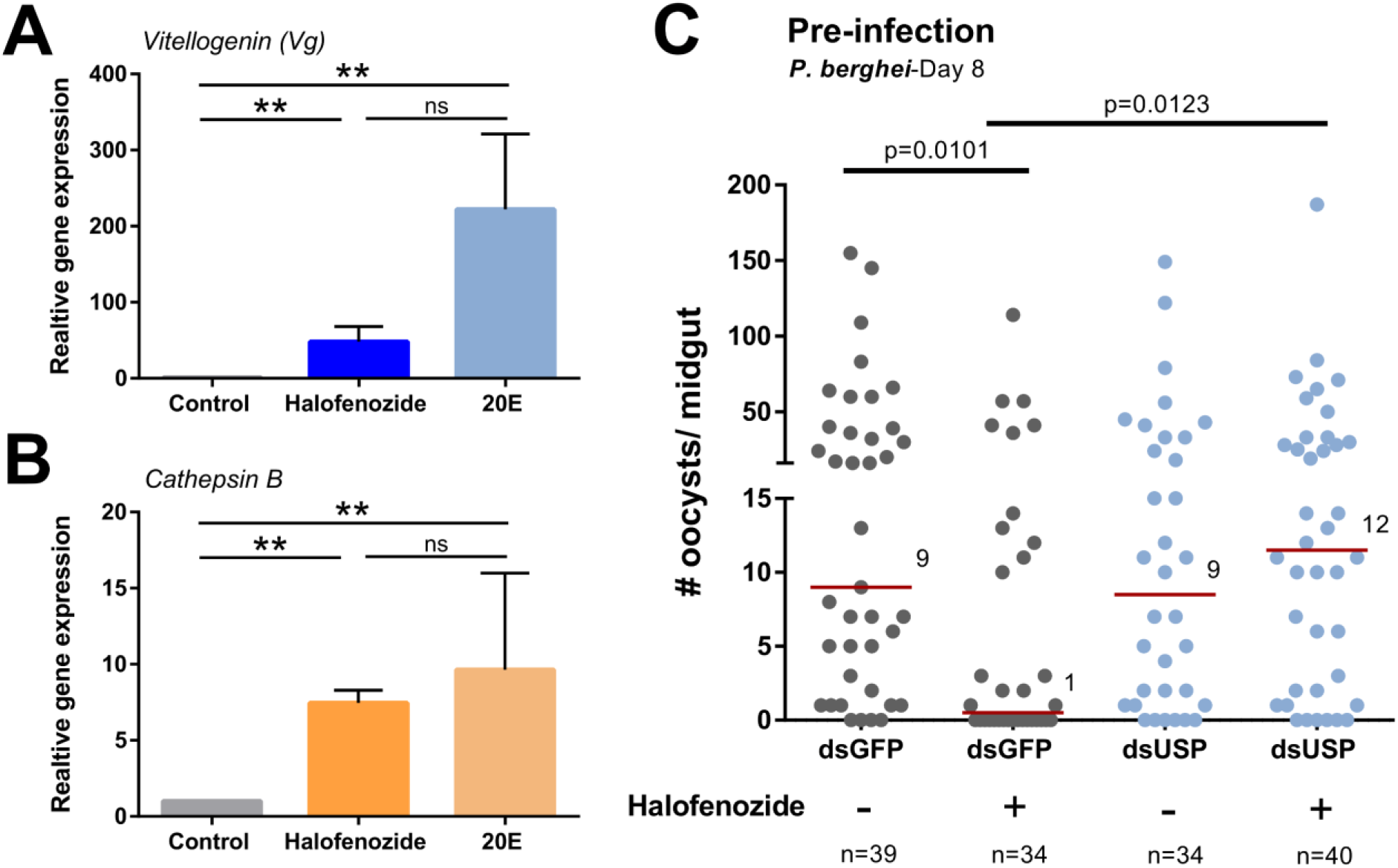
Halofenozide stimulates ecdysone signaling and requires ultraspiracle (USP) to confer anti-*Plasmodium* immune priming. The effects of halofenozide (0.5 μg/μL), application on mosquito gene expression were evaluated on two downstream components of ecdysone signaling, *vitellogenin* **(A)** and *cathepsin B* **(B)**, and compared to 20E as a positive control. Statistical significance was determined by Mann-Whitney analysis (**, *P*< 0.01; ns, not significant) from three or more independent biological samples. **(C)** To determine the involvement of the heterodimeric EcR/USP receptor in mediating the effects of halofenozide priming, we examined *USP* function by RNAi in *GFP* (control)*-* or *USP-*silenced mosquitoes. Two days post dsRNA injection, we topically applied acetone (-, control) or halofenozide (+, 0.5 μg/μL) and challenged mosquitoes with *P. berghei* ~24 hours post-halofenozide topical application. Oocyst numbers were evaluated eight days for parasite development. The red bar delineates the median number of oocysts from pooled data from three independent experiments. Data were analyzed by Mann-Whitney analysis using GraphPad Prism 6.0. n= the number of mosquitoes examined for each condition.

### Mechanisms of halofenozide immune induction are distinct from 20E

We previously examined the influence of 20E on cellular immunity and the effects of 20E priming that limit malaria parasite infection ^8^. Similarly, we wanted to determine if halofenozide application promotes similar changes to 20E in gene expression, cellular immunity, and anti-microbial defense. To compare gene expression, we examined the expression of *cecropin 1* (*CEC 1*) and *cecropin 3* (*CEC 3*), which previously were significantly up-regulated in response to 20E injection in naïve mosquitoes ^8^. However, halofenozide did not stimulate the expression of *CEC1* or *CEC3* (Figure 5A and 5B), suggesting that halofenozide initiates a different transcriptional repertoire than that of 20E^8^. This led us to question whether halofenozide application primed cellular immunity similar to 20E in naïve mosquitoes ^8^. Contrary to previous work with 20E, halofenozide treatment did not influence the phagocytic activity of mosquito immune cells (Figure 5C). Together, these results suggest that the mechanisms of halofenozide immune induction are distinct from that of 20E in limiting malaria parasites.

**Figure 5.**
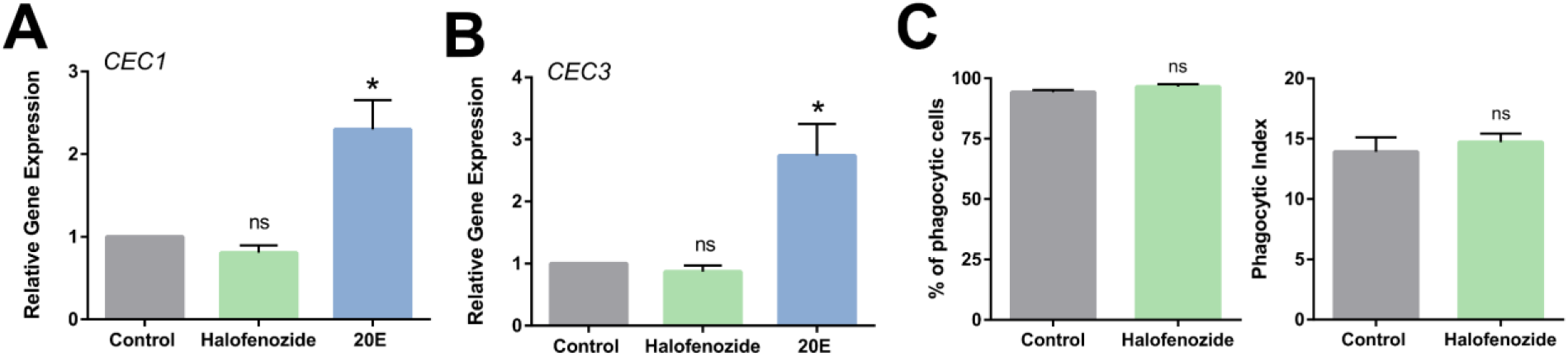
Halofenozide does not influence AMP production or cellular immunity. The effects of halofenozide topical application and 20E injection were compared by examining their respective influence on gene expression using the 20E-responsive anti-microbial peptides (AMPs), cecropin 1 (CEC1) **(A)** and cecropin 3 (CEC3) **(B)**. Statistical significance was determined by Mann-Whitney analysis (*, *P*<0.05) from three or more independent biological samples. The influence of halofenozide was also examined on the phagocytic activity of mosquito immune cells, evaluating the percentage of phagocytic cells and the phagocytic index (number of beads per cell) **(C)**. Data was analyzed by Mann-Whitney analysis using GraphPad Prism 6.0 to determine significance. ns, not significant.

## Discussion

To date, the most effective malaria control strategies target the mosquito host by reducing mosquito habitats, preventing transmission through the use of bed nets, or by killing adult mosquitoes through the use of insecticides ^20^. However, due to increasing insecticide resistance in many malaria-endemic regions of the world, it is critical to develop new methods or improve on existing tools to interrupt malaria transmission. This includes the use of commonly used insecticides to manipulate mosquito host physiology, influencing fitness or rendering them less likely to acquire and transmit mosquito-borne pathogens. Childs *et al.* demonstrated that the application of a 20E agonist, DBH, negatively influenced mosquito survival, reproduction, and reduced *P. falciparum* survival in the mosquito host ^10^, supporting that 20E agonists have the potential to improve existing malaria control strategies ^10,21^. This is further supported by recent studies demonstrating that 20E confers anti-*Plasmodium* immunity ^8^, yet prior to our presented work herein, the manner in which 20E agonists influence *Plasmodium* survival have remained unknown.

In agreement with previous work with DBH ^10^, we found the topical application of halofenozide, a similar 20E agonist, significantly impaired *P. berghei* oocyst survival and reduced infection prevalence. Additionally, we provide new details into the temporal effects of halofenozide immune activation, demonstrating that the anti-*Plasmodium* properties of halofenozide treatment only function to reduce *Plasmodium* numbers when applied prior to malaria parasite challenge. Therefore, if 20E agonists such as halofenozide are integrated into bed nets or employed by indoor residual spraying (IRS) as previously proposed ^10,21^, mosquitoes must to come into contact prior to taking an infectious blood meal to influence parasite survival. However, further studies are required to determine if other 20E agonists, such as DBH, share similar mechanisms of limiting malaria parasites.

Several lines of evidence suggest that halofenozide application promotes “early-phase” components of anti-*Plasmodium* immunity that influence the success of ookinete invasion ^5,16,22^. This includes the reduction in the number of invading ookinetes, as well as early oocyst number. These data are further supported by our results demonstrating that halofenozide application ~24h post-infection, and after ookinete invasion, did not influence malaria parasite survival. As a result, halofenozide (and potentially other 20E agonists) may influence complement function ^23–25^, or a yet undescribed mechanism to limit *Plasmodium* ookinete survival. However, further experiments are required to define the direct mechanisms by which halofenozide promotes malaria parasite killing.

The integral role of USP in conferring the effects of halofenozide application provide strong support that halofenozide activates ecdysone signaling, which is in agreement with previous evidence that 20E agonists competitively bind to the EcR/USP heterodimer ^26^. Although we were unable to knock-down *EcR*, EcR alone is incapable of high affinity binding to 20E to promote the activation of downstream 20E-regulated genes ^27^. This is further supported by the activation of *Vg* and *cathepsin B,* two highly responsive 20E-regulated genes ^19,28^, following halofenozide application at comparable levels to the injection of 20E. Therefore, our data support that halofenozide-mediated immune induction occurs through the heterodimeric ecdysone receptor.

Previous work has identified the downstream targets of 20E signaling in *An. gambiae*, as well as demonstrated the influence of 20E on cellular immunity and anti-pathogen defense responses that limit bacterial and parasite survival ^8^. Since halofenozide functions through the canonical 20E receptor, we originally hypothesized that halofenozide and 20E would similarly influence mosquito physiology and immunity ^8^. While both 20E and halofenozide stimulate *Vg* and *cathepsin B*, halofenozide application does not stimulate immune gene expression or cellular immunity similar to 20E ^8^. Together, these results suggest that halofenozide and 20E work through similar, yet distinct mechanisms to influence mosquito physiology and promote anti-*Plasmodium* immunity. This may potentially be explained by differences between steroid hormones such as 20E and their non-steroidal agonist to enter target cells and stimulate cellular function ^29^. In support of this hypothesis, previous work in *Drosophila* demonstrates that 20E and chromafenozide, a nonsteroidal 20E agonist, differentially enter cells to stimulate cell function ^12^. 20E requires the ecdysone importer (*EcI*) to gain cell entry, while chromafenozide can stimulate cellular responses independent of EcI ^12^. Based on these notable differences between steroid hormones and non-steroidal agonists, additional studies are required to further delineate the mechanisms of 20E and halofenozide immune induction in the mosquito host.

In summary, we demonstrate that the 20E agonist, halofenozide, is an effective tool to promote physiological responses that reduce the prevalence and intensity of malaria parasite infection, similar to previous studies with the 20E agonist DBH ^10^. Moreover, we define the temporal requirements of halofenozide application, where *Plasmodium* infection intensity is limited only in mosquitoes treated pre-infection, consistent with a role in halofenozide mediating physiological responses that limit the success of ookinete invasion. We demonstrate that halofenozide requires the function of USP, implicating ecdysone signaling in mediating the physiological responses that limit *Plasmodium* infection. However, through comparative analysis of gene expression and cellular immune function, we establish that halofenozide application produces physiological responses not directly comparable to the effects of 20E ^8^. From these data, we provide new fundamental insight into the mechanisms of halofenozide immune induction to better understand 20E agonist function. Together, the evidence presented here and in previous studies ^10,21^ suggest that 20E agonists could be an effective tool to help reduce the transmission of malaria.

## Methods

### Ethics statement

The protocols and procedures used in this study were approved by the Animal Care and Use Committee at Iowa State University (IACUC-18-228). All experiments were performed in accordance with the relevant guidelines and regulations of the approved study.

### Mosquito rearing and *Plasmodium* infections

*An. gambiae* (G3 strain) were maintained at 27°C and 80% relative humidity, with a 14/10-hour light/dark cycle. Larvae were maintained on a ground fish food diet (Tetramin). Pupae were isolated using a pupal separator (John W. Hock Company) and placed in containers of ~50 mosquitoes where they were allowed to eclose in mixed populations of male and female mosquitoes. Adult mosquitoes were maintained on a 10% sucrose solution.

For mosquito infections with *P. berghei*, a mCherry *P. berghei* strain ^30^ was passaged into female Swiss Webster mice as previously described ^8,16,31,32^. After the confirmation of an active infection by the presence of exflagellation, mice were anesthetized and placed on mosquito cages for feeding. Infected mosquitoes were sorted and then maintained at 19°C. Mosquito midguts were dissected eight days post-infection in 1x PBS, and oocyst numbers were measured by fluorescence using a Nikon Eclipse 50i microscope.

### Halofenozide topical application on adult mosquitoes

For topical applications with halofenozide (Chem Services Inc.), a stock solution of 40μg/μL was prepared using 0.04 grams of halofenozide in 1 mL of 100% acetone. Prior to application, working concentration were prepared by diluting the stock solution 1:79 or 1:159 in 100% acetone to achieve respective working concentrations of 0.5 μg/μL and 0.25μg/μL. Adult female *An. gambiae* (3-5 days post eclosion) were topically applied with 0.2μL of either 100% acetone (control) or halofenozide (0.25 μg/μL and 0.5μg/μL) in 100% acetone. Applications were performed on individual mosquitoes using a repeating syringe dispenser (PB600-1 Hamilton syringe with a 10-microliter syringe). Surviving mosquitoes were challenged (*E. coli* or *P. berghei*) 24 hours post-topical application or were used to collect samples for RNA isolation to determine the effects of halofenozide treatment on gene expression.

### Gene expression analysis

Total RNA was isolated from whole mosquitoes (~10 mosquitoes) using TRIzol (Thermo Fisher Scientific) according to the manufacturer’s protocol. RNA samples were quantified using a Nanodrop spectrophotometer (Thermo Fisher Scientific) and ~2μg of total RNA was used as a template for cDNA synthesis using the RevertAid First Strand cDNA Synthesis kit (Thermo Fisher). Gene expression was measured by qRT-PCR using gene-specific primers (Table S1) and PowerSYBR Green (Invitrogen). Samples were run in triplicate and normalized to an S7 reference gene and quantified using the 2^−ΔΔCT^ method as previously ^33^.

### dsRNA synthesis and gene-silencing

T7 primers for GFP and *ultraspiricle* (USP) were used to amplify cDNA prepared from whole *An. gambiae* mosquitoes and cloned into a pJET1.2 vector using a CloneJET PCR Cloning Kit (Thermo Fisher). The resulting plasmids were used as a template for amplification using the corresponding T7 primers (Table S2). PCR products were purified using the DNA Clean and Concentrator kit (Zymo Research) and used as a template for dsRNA synthesis as previously described ^8,31^. The resulting dsRNA was resuspended in RNase-free water to a concentration of 3 μg/uL. For gene-silencing experiments, 69nL of dsRNA was injected per mosquito and evaluated by qRT-PCR to establish gene knockdowns at 1-5 days post-injection. Time points with the highest efficiency of gene-silencing were chosen for downstream experiments.

### Phagocytosis Assays

Phagocytosis assays were performed *in vivo* as previously described ^8,32^ using red fluorescent FluoSpheres (1 μm Molecular Probes) at a 1:10 dilution in 1x PBS. Following injection, mosquitoes were allowed to recover for two hours at 19°C, then perfused onto a multi-test slide. To visualize hemocytes, cells were stained using a 1:500 dilution of FITC-labeled wheat germ agglutinin (WGA; Sigma) in 1x PBS, while nuclei were stained with ProLong Gold anti-fade reagent with DAPI (Invitrogen). Hemocytes were identified by the presence of WGA and DAPI signals, with the number of cells containing beads by the total number of cells to calculate the percent phagocytosis. The phagocytic index was calculated by counting the total number of beads per cell (this is summed for all of the cells) dividing by the number of phagocytic cells. Approximately 200 cells were counted per mosquito sample.

### Evaluation of halofenozide impacts on bacteria

The potential impacts of halofenozide on the mosquito microbiota were evaluated using both *in vivo* and *in vitro* experiments. For *in vivo* experiments, we analyzed the relative quantification of 16s rRNA expression ^15,16^ as a proxy for mosquito microbiota titers between acetone (control) and halofenozide (0.5μg/μL) treatments. cDNA was prepared from whole mosquitoes receiving topical applications as described above, with 16s rRNA expression examined by qRT-PCR using universal 16S bacterial primers listed in Table S1. For *in vitro* experiments, *E. coli* was cultured in Luria Bertani (LB) broth over night at 37°C, then used to seed a bacterial suspension (OD_600_=0.4) in 2 ml of LB broth. To determine the potential impacts of halofenozide on bacterial growth, 2 μl of either halofenozide (0.25 μg/μl and 0.5 μg/μl) or 100% acetone were added to bacterial suspensions and continually cultured at 37°C with shaking at 210 rpm. Bacterial cultures were measured by optical density (OD) at 600 nM every 2 h up to 6 h post-challenge to monitor bacterial growth.

### Mosquito ookinete invasion assay

The influence of halofenozide on *Plasmodium* sexual development and ookinete invasion was examined *in vivo* as previously described ^34^. Mosquitoes were topically treated with acetone and halofenozide (0.5 μg/μl) as described above, then challenged with *P. berghei* infection. At 18 h post infection, mosquito midguts were dissected dissociating the blood bolus and midgut. For each mosquito, the proportion of each parasite developmental stage (zygote or ookinete) was determined by counting approximately 50 parasites from random chosen fields. In addition, the number of live ookinetes (as determined by mCherry fluorescence of the transgenic *P. berghei* strain) was determined for each of the dissociated midgut samples to assess parasite invasion.

### Ookinete culture

*In vitro* culture of *P. berghei* ookinetes were performed as previously described ^35^. Briefly, mice were injected i.p. with 200 μL of 10 mg/mL phenylhydrazine/1× PBS (Sigma). Three days later, mice were infected i.p. with 10^8^*P. berghei*-GFP iRBCs. Three days later, the *P. berghei* gametocyte-enriched blood was collected by cardiac puncture and cultured in 25 ml flasks with ookinete culture medium containing 0.5μg/μl of halofenozide in 100% acetone (1:1000 dilution) or an equal volume of 100% acetone, for 24 h at 19 °C with gentle agitation. Parasites were fixed with 4% w/v paraformaldehyde and stained with an anti-Pbs21 antibody ^36^. Pbs21-positive cells including gametes and zygotes (round forms), retorts and ookinetes were counted under a fluorescence microscope.

### Ookinete motility assays

*P. berghei*-GFP ookinetes cultured in the presence of 0.5μg/μl of halofenozide in 100% acetone (1:1000 dilution) or an equal volume of 100% acetone were mixed with an equal volume of Geltrex LDEV-Free Reduced Growth Factor Basement Membrane Matrix (Thermo Fisher Scientific). The mixture was applied onto a glass slide, then covered with a round coverslip and incubated at room temperature for 5 min to allow the matrix to solidify. Ookinete motility was immediately monitored for 15 min at room temperature with frames taken every 15 s on an Axio Imager M2 fluorescence microscope using a 20x objective and an Axiocam 506 mono camera (Zeiss). Videos were acquired and processed using the Zen 2.5 Software (Zeiss).

## Acknowledgments

We would like to acknowledge David Hall for his efforts in mosquito rearing. This work was supported by a National Science Foundation Graduate Research Fellowship Program under Grant No. 1744592 (R.A.R), the NIH Distinguished Scholars Program and the Intramural Research Program of the Division of Intramural Research AI001250-01, NIAID, National Institutes of Health (J. V-R.), and by support from the Agricultural Experiment Station at Iowa State University to (R.C.S.).

