## Supplementary material for "The 20-hydroxyecdysone agonist, halofenozide, promotes anti-*Plasmodium* immunity in *Anopheles gambiae* via the ecdysone receptor": Figures S1-S4, Tables S1-S2

### Supplemental Information

#### Supplemental Figure Legends

**Figure S1.** Mosquito survival following halofenozide application.

**Figure S2.** Halofenozide does not influence bacteria.

**Figure S3.** Increasing concentrations of halofenozide stimulate higher levels of ecdysone signaling.

**Figure S4.** Targeting the heterodimeric ecdysone receptor (EcR/USP) by RNAi.

#### Supplemental Tables

**Table S1.** Primers for qRT-PCR analysis.

**Table S2.** Primers for dsRNA synthesis.=

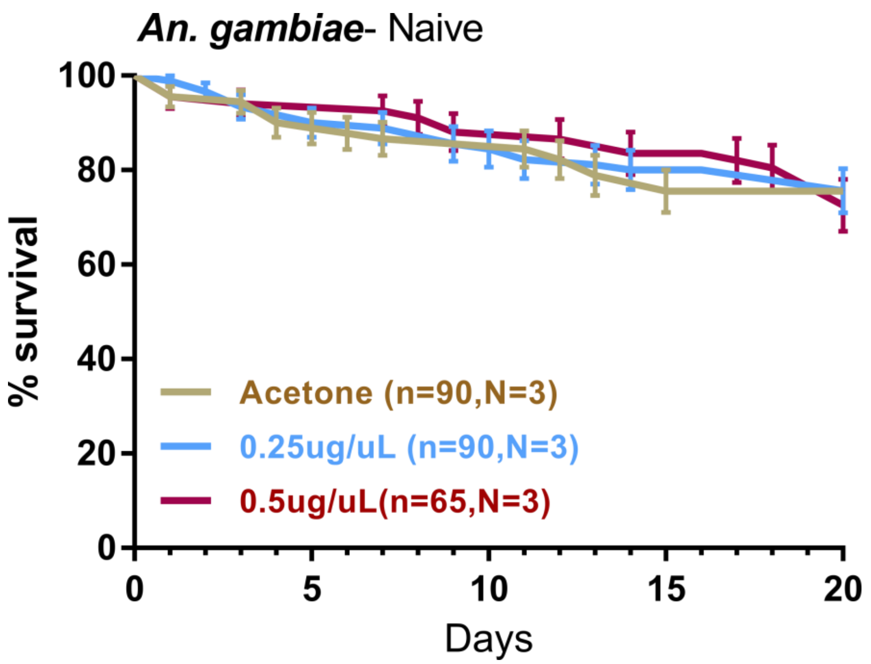

**Figure S1. Mosquito survival following halofenozide application.** Naive female *An. gambiae* mosquitoes were topically applied with either acetone (control) or halofenozide (0.25 µg/µL and 0.5 µg/µL). Mosquito survival was monitored for 20 days post-application. No significant differences between treatments were detected when a Log-rank (Mantel-Cox) test was performed.

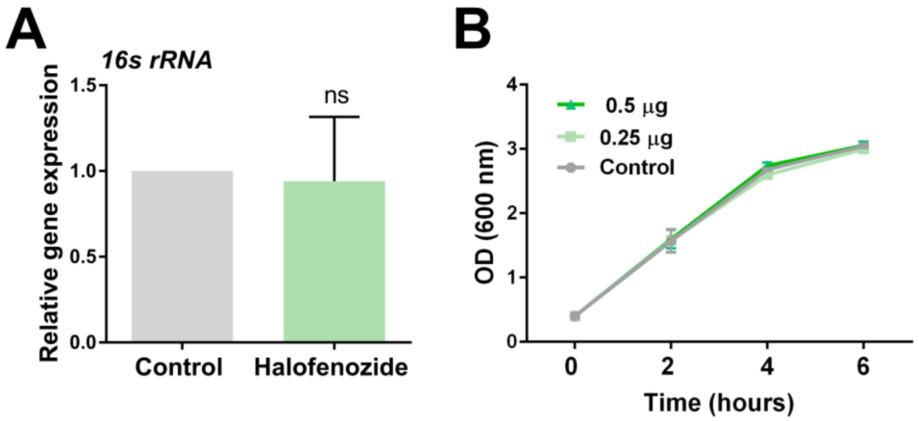

**Figure S2. Halofenozide does not influence bacteria.** The influence of halofenozide on the mosquito microbiota was examined in control- or halofenozide-treated (0.5 μg) mosquitoes using 16s rRNA primers by qRT-PCR (**A**). Results display previously results from four independent experiments. Data were examined using Mann-Whitney analysis. ns, not significant. Bacteria growth was also examined *in vitro*, in which halofenozide (0.25 and 0.5 μg/ml) or acetone (control) were added to liquid cultures of *E. coli* and their growth was examined for 6 hours using measurements of optical density (OD) at 600 nm (**B**). Experiments were performed in two independent experiments and analyzed using a two-way ANOVA and Bonferroni's multiple comparison test. No significant differences were detected between treatments at any experimental timepoint.

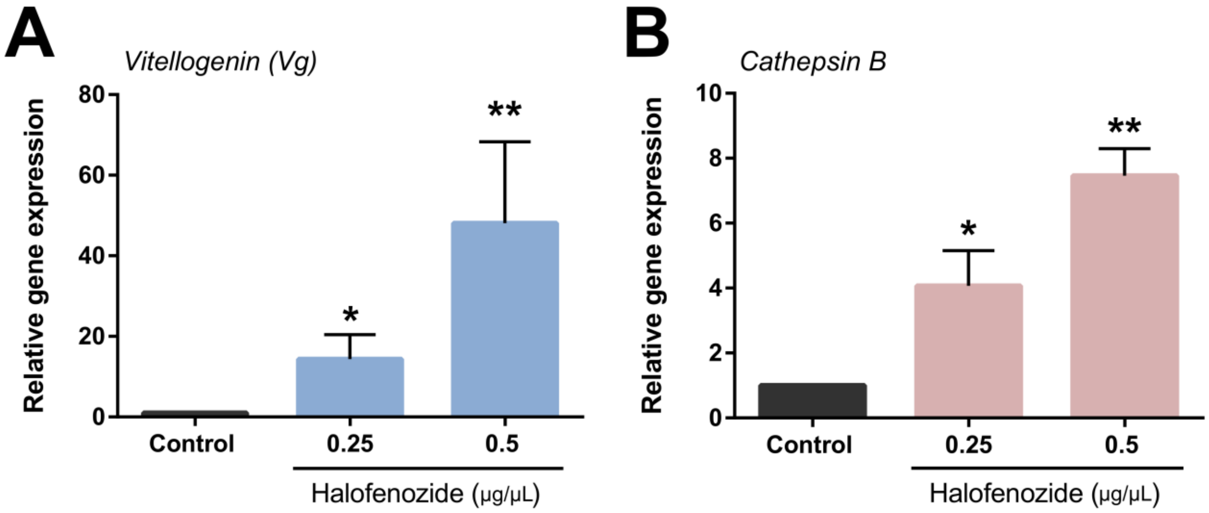

**Figure S3. Increasing concentrations of halofenozide stimulate higher levels of ecdysone signaling.** Analysis of vitellogenin (A) and cathepsin B (B) gene expression by qRT-PCR in response to increasing concentrations of halofenozide in whole naive female *An. gambiae*. Data from three or more experiments were compared to control (acetone) and analyzed using a Mann-Whitney test in GraphPad Prism 6.0 to determine significance (*, $P<0.05$; **, $P<0.01$).

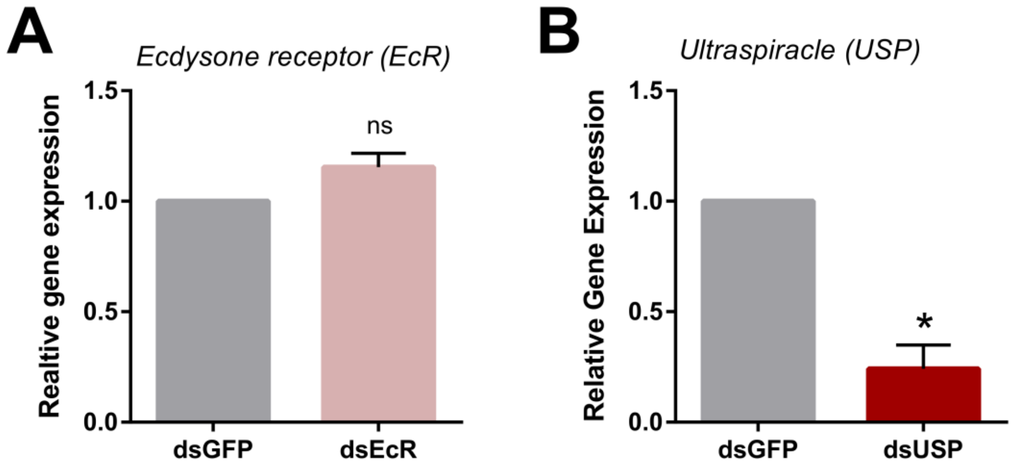

**Figure S4. Targeting the heterodimeric ecdysone receptor (EcR/USP) by RNAi.** To disrupt the heterodimeric ecdysone receptor (EcR/USP), EcR and USP were individually targeted by the injection of specific dsRNA. To validate the presence or absence of a knockdown, the expression of *EcR* **(A)** and *USP* **(B)** in whole mosquitoes 2 days post-dsRNA injection by qRT-PCR. Data from three or more experiments were analyzed using a Mann-Whitney test in GraphPad Prism 6.0 to determine significance (*, $P<0.05$).

**Table S1. Primers for qRT-PCR analysis**

| Primer | Gene ID | Sequence (5'- 3') |
|---|---|---|
| Cathepsin B-F | AGAP004534 | GCCAACGGTCTAGTGTCGAGTGTCG |
| Cathepsin B-R | | ACTCGTACCGTCTGATCGGCACCTT |
| Cecropin 1-F | AGAP000693 | TTCATCTTTGTCGTGCTGGC |
| Cecropin 1-R | | GCACTGCCAGCACGACAAAG |
| Cecropin 3-F | AGAP000694 | ACGTACTGAACCACCTGCGCGTT |
| Cecropin 3-R | | GCGCTGTGTGCGCCGATGAA |
| rpS7-F | AGAP010592 | ACCCCATCGAACACAAAGTTGACACT |
| rpS7-R | | CTCCGATCTTTCACATTCCAGTAGCAC |
| USP-F | AGAP002095 | TGAAGTCCGAAGAAATCAACTCGAC |
| USP-R | | GGGCAAACTCGATTAGCTGGTAGAT |
| Vg-F | AGAP004203 | TGCAGTACATCGAGCAGGGTGACAA |
| Vg-R | | CTTGACGGTCTTGGTGACCGACTTG |
| Universal bacteria 16S-F | N/A | TCCTACGGGAGGCAGCAGT |
| Universal bacteria 16S-R | | GGACTACCAGGGTATCTAATCCTGTT |

**Table S2. Primers for dsRNA synthesis**

| Primer | Gene ID | Sequence (5'- 3') |
|---|---|---|
| GFP T7-F | | TAATACGACTCACTATAGGGAGAATGGTGAGCAAGGGCGAGGAGCTGT |
| GFP T7-R | | TAATACGACTCACTATAGGGAGATTACTTGTACAGCTCGTCCATGCC |
| USP T7-F | AGAP002095 | TAATACGACTCACTATAGGGCCTAAAATGTCG |
| USP T7-R | | TAATACGACTCACTATAGGGATCGGGAGACACACGCAGT |
